# Identification and characterization of *Aedes albopictus* long noncoding RNAs provides insights into their roles in development and flavivirus infection

**DOI:** 10.1101/448852

**Authors:** Azali Azlan, Muhammad Amir Yunus, Ghows Azzam

## Abstract

*Aedes albopictus* (*Ae. albopictus*) is an important vector of arboviruses such as Dengue virus (DENV), Chikungunya virus (CHIKV), and Zika virus (ZIKV). Long noncoding RNA (lncRNAs) have been identified in other vectors including *Aedes aegypti* and *Anopheles* mosquitoes, few of which have been implicated in immunity and viral replication. To identify lncRNAs with potential biological functions in *Ae. albopictus*, we performed RNA-seq on *Ae. albopictus* cells infected with DENV and ZIKV, and analyzed them together with public datasets. We identified a total of 23,899 transcripts, 16,089 were intergenic while 3,126 and 4,183 of them were antisense and intronic to annotated genes respectively. *Ae. albopictus* lncRNAs shared many of the characteristics with their invertebrate and vertebrate counterparts, such as low expression, low GC content, short in length, and low conservation even among closely related species. Compared to protein-coding genes, lncRNAs exhibited higher tendency to be expressed in a stage-specific manner. Besides, expression of lncRNAs and nearest protein-coding genes tended to be correlated, especially for the gene pairs within 1kb from each other. We also discovered that *Ae. albopictus* lncRNAs have the potential to act as precursors for miRNA and piRNAs, both of which have been implicated in antiviral defense in *Aedes* mosquito. Upon flavivirus infection, lncRNAs were observed to be differentially expressed, which possibly indicates the involvement of lncRNAs in the host-antiviral defense. Our study provides the first systematic identification of lncRNAs in *Ae. albopictus*, hence, offering a foundation for future studies of lncRNA functions.

## Introduction

The Asian tiger mosquito, *Aedes albopictus* (*Ae. albopictus*) is an important vector of arboviruses such as Dengue virus (DENV), Chikungunya virus (CHIKV), and Zika virus (ZIKV). Due to its invasiveness and aggressive spread, *Ae. albopictus* has widespread geographic distribution, posing serious health threat across the globe in both tropical and temperate regions. Although *Ae. albopictus* is considered as less competent vector of DENV than *Aedes aegypti* (*Ae. aegypti*), *Ae. albopictus* was responsible for dengue outbreaks in Hawaii, China, and Europe, primarily because of its fast expansion across the globe (Chen et al., 2015).

Although protein-coding genes have been the central focus, many reports have indicated that noncoding RNAs (ncRNAs), such as long noncoding RNAs (lncRNAs) and small RNAs, play important roles in development and virus-host interaction (Etebari et al., 2017, 2016; Liu et al., 2015; Miesen et al., 2016a, 2016b). Although lncRNAs lack coding potential, similar to mRNAs, they are the products of Pol II, and they undergo polyadenylation, capping and alternative splicing (Ulitsky and Bartel, 2013). Due to their mRNA-like features, lncRNAs are usually represented in RNA-seq datasets. Next-generation sequencing allows quick genome-wide identification of lncRNAs including the lowly expressed transcripts, and this technology is independent on complete genome and gene annotation, making it an ideal strategy to detect novel lncRNAs (Wang et al., 2009; Wilhelm et al., 2010).

lncRNAs act by various mechanisms. Several lncRNAs have been shown to modulate the chromatin state, thereby; regulating gene expression inside cells (Wang et al., 2011) Meanwhile, other lncRNAs were associated with post-transcriptional regulation such as Malat1, which is important for alternative splicing of mRNA transcripts (Tripathi et al., 2010). lncRNA was also shown to be involved in virus-host interaction in *Ae. aegypti*. For example, knockdown of lincRNA_1317 in *Ae. aegypti* cells resulted in an increased replication of DENV (Etebari et al., 2016). Another potential role of lncRNAs is to act as precursors or templates for the generation of mature small RNAs (Yoon et al., 2014). Small RNAs in metazoa can be categorized into three distinct groups based on their biogenesis and mechanism of action (Azlan et al., 2016; Siomi et al., 2011): microRNAs (miRNAs), small-interfering RNAs (siRNAs), and PIWI-interacting RNAs (piRNAs). miRNAs, ~22 nucleotide (nt) in length, function in the regulation of gene expression in both animals and plants by targeting cognate messenger RNAs (mRNAs) via imperfect base-pairing, resulting in either mRNA cleavage or translational repression (Mallory and Vaucheret, 2010). siRNAs are ~ 21 to ~24 nt RNAs in length that originate from long double-stranded RNA (dsRNA) or hairpins, both of which can be encoded endogenously in the genome or can be exogenously introduced into the cells. piRNAs (~24-35 nt in length) function to control the activity of transposable element (TE); thereby, ensuring the inheritance of genomic information from one generation to another unscathed (Siomi et al., 2011). Studies of piRNAs and their protein partners, PIWI proteins, in mice and flies have led to the proposal of two piRNA biogenesis pathways: primary pathway and secondary ping-pong amplification cycle (Brennecke et al., 2007). piRNA biogenesis begins with the transcription of piRNA precursor mainly from distinct loci in the genome that are referred to as piRNA clusters. The primary biogenesis pathway is thought to contribute to the initial population of piRNAs, and the ping-pong amplification cycle, then amplifies the piRNA pool that targets active TE. The two pathways work together to elicit effective defense against active transposons. (Brennecke et al., 2007; Siomi et al., 2011).

Here, we report the first systematic genome-wide identification and characterization of lncRNAs in *Ae. albopictus*. To identify a set of high confident lncRNA transcripts, we performed RNAseq based de novo transcript discovery, and applied stringent filtering of transcripts having coding potential. We then characterized each lnRNA by many features such as transcript structures, conservation, and developmental expression. We also investigate the functional link of lncRNAs and small RNAs, especially miRNAs and piRNAs. Beside that, we examined the lncRNA expression landscape upon viral-infection to gain insights into the lncRNA functions in virus-host interaction. Although our knowledge on mosquito-virus interaction in *Ae. albopictus* is still limited, results generated from this study will provide invaluable resources for future investigations. Thorough understanding on the intricate relationship between viruses and mosquitoes at cellular level, which involved both coding and non-coding genes, is crucial for devising efficient strategies for vector control.

## Results

### Genome-wide identification of lncRNAs in *Ae. albopictus*

To identify novel lncRNAs, we generated 9 paired-end RNA-seq libraries (triplicate of C6/36 cells at rest, DENV1-infected, and ZIKV-infected) and analyzed them together with 185 publicly available RNA-seq libraries generated from *Ae. albopictus*. To get high confident lncRNA transcripts, we applied a stringent identification pipeline that was adapted with slight alterations from lncRNA identification studies in other species (Azlan et al., 2018; Chen et al., 2016; Etebari et al., 2016; Hezroni et al., 2015; Wu et al., 2016; Young et al., 2012). An overview of lncRNA identification pipeline can be found in **Figure 1.** The pipeline began with alignment of each RNA-seq library against *Ae. albopictus* genome (assembly: canu_80X_arrow2.2, strain: C6/36, VectorBase) using HISAT2 (Kim et al., 2015), followed by assembly of RNA-seq alignments into potential transcripts by Stringtie (Pertea et al., 2015). The transcript assemblies were then merged using Stringtie, yielding a total of 252,453 transcripts derived from 137,743 loci.

**Figure 1.**
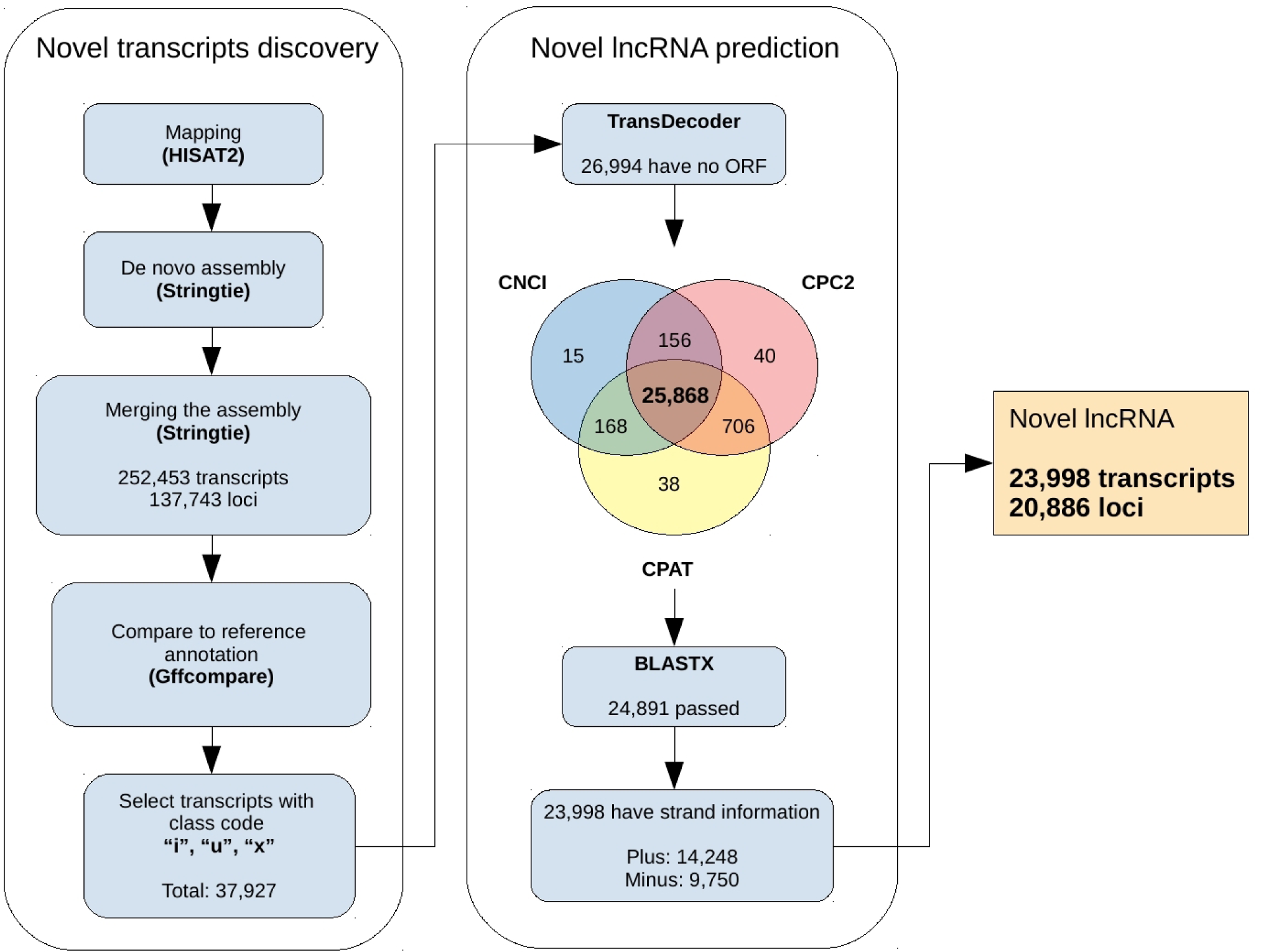
lncRNA identification pipeline

We annotated and compared the transcripts with reference annotation using Gffcompare (Trapnell et al., 2010). For downstream analysis, we only chose novel transcripts that were intergenic, intronic and antisense to the reference genes – all of which made a total of 37,927 transcripts. Out of 37,927 transcripts, 10,933 were shown to be coding by TransDecoder, and they were discarded. The remaining 26,994 transcripts were subjected for coding potential assessment by three different softwares, namely CPAT (Wang et al., 2013), CPC2 (Kang et al., 2017), and CNCI (Sun et al., 2013). We found that 25,868 transcripts were identified to be noncoding in all three algorithms. We then performed BLASTX against Swissprot protein database, and 977 transcripts having significant hits (E-value < 10-5) were discarded. Finally, we examined the strandedness of the remaining 24,891 transcripts, and transcripts without strand information were removed. Detailed description on the prediction analysis and parameter used can be found in **Material and Methods** section.

In this study, we identified a set of 23,998 novel lncRNA transcripts derived from 20,886 loci. The number of intergenic, intronic and antisense lncRNAs were 16,089, 4,183, and 3,126 respectively. Current annotation listed 8,571 lncRNA transcripts; hence, altogether, the total number of lncRNAs in *Ae. albopictus* were 32,569 transcripts.

### *Ae. albopictus* lncRNAs shared similar genomic features with other species

Studies done in other species revealed that, compared to protein-coding gene, lncRNAs are typically shorter in length, have low GC content, have high repeat contents, and their sequence conservation is relatively low even among closely related species (Azlan et al., 2018; Chen et al., 2016; Etebari et al., 2016; A. Pauli et al., 2012; Wu et al., 2016; Young et al., 2012). Our analysis showed that *Ae. albopictus* lncRNAs shared similar characteristics with their vertebrate and invertebrate couterparts. We found that lncRNA transcripts were shorter than protein-coding mRNA (**Figure 2A)**. Coding mRNA transcripts had a mean length of 2,659 bp, while the average size of novel and known lncRNAs was 662.9 bp and 697 bp respectively. Besides, both novel and known lncRNAs had lower GC content than the coding transcripts. Average GC content of novel and known lncRNAs were 42.4% and 42.8% respectively while coding sequence had the mean GC content of 51.3% (**Figure 2B)**. Not only lncRNAs had lower GC content, but other non protein-coding sequence in the genome, including intron, intergenic regions, 5’UTR and 3’UTR, also had lower GC content.

**Figure 2.**
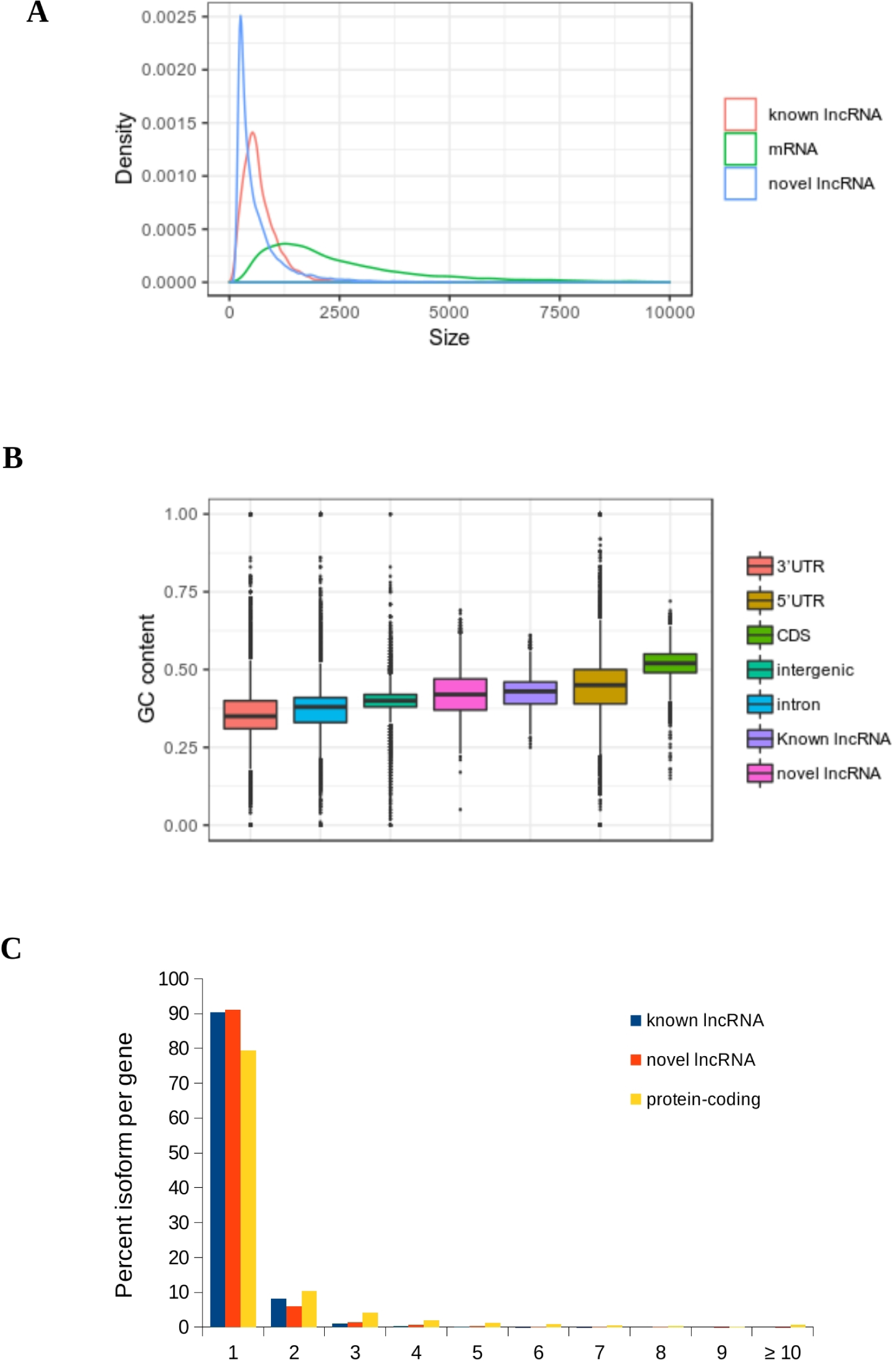
Characterization of *Ae. albopictus* lncRNA (A) Size distribution of mRNA transcripts, novel and known lncRNA (B) GC content (C) Number of isoform per gene

Similar to previous reports (Hezroni et al., 2015; Nam and Bartel, 2012; Wu et al., 2016), we found that lncRNA transcripts had higher composition of repeat content compared to coding mRNA 2.37% and 0.08% on lncRNA and protein-coding exon nts respectively were embedded within repeat elements. Repeat elements embedded within lncRNAs include hellitron, LINE, SINE, satellites, and LTR. More than 90% of both novel and known lncRNAs had one isoform per gene, and lncRNA genes having more than 10 isoforms were less than 10 (**Figure 2C)**. lncRNA loci possesed a slightly lower isoforms per gene than protein-coding gene. lncRNA had an average of 1.14 isoforms per gene locus, while protein-coding harbored 1.5 transcripts per gene. Shorter length, lower GC content, and less number of isoform per genes suggest that lncRNAs are less complex than protein-coding transcripts.

To examine the level of sequence conservation, we performed BLASTN of lncRNA transcripts against the genomes of closely related insects species namely *Ae. aegypti, C, quinquifasciatus*, *An, gambiae* and *D. melanogaster*. From BLASTN results, we defined conserved lncRNAs as transcripts having E-value less than 10^-50^ (Etebari et al., 2016). Only 19 lncRNA transcripts were conserved in all 4 genomes being tested, and most of the conserved lncRNAs (5,224 transcripts) shared high shared sequence similarity with *Ae. aegypti* genome. On the contrary, coding mRNAs showed higher level of sequence conservation by BLASTN (E-value < 10^-50^) method, as shown by the number of conserved transcripts shared in all genomes and the number of transcripts shared between *Ae. aegypti* (**Supplemental Figure 2**). Overall, total number of lncRNA transcripts that shared high sequence similarity with *Ae. aegypti* was significantly lower than that observed in mRNAs. For instance, only 16% (5,224 out of 32,569 transcripts) of lncRNAs were highly conserved, but for coding mRNAs, 78% of their total transcripts (33,431 of 42,899 transcripts) exhibited high level of sequence conservation.

### *Ae. albopictus* lncRNAs may act as precursors for miRNAs and piRNAs

In verterbrates, some lncRNAs were processed to generate miRNAs, showing that lncRNA transcripts act as precursors in miRNA biogenesis (Yoon et al., 2014; Zhang et al., 2018, 2017). To examine if this was also the case for *Ae. albopitus* lncRNAs, we examined lncRNA genomic coordinates that were fully overlapped with miRNA precusor loci. Due to the fact that miRNA annotation was not systematically done in *Ae. albopictus* C6/36 genome (canu_80X_arrow2.2 assembly), we sought to produce comprehensive list of miRNAs by analyzing 20 public small RNA datasets (accession: SRA060684 and SRP096579) using miRDeep2 software (Friedländer et al., 2012). We predicted a total of 116 and 10 known and novel miRNAs (**Supplemental Table 2**). We defined known miRNAs as those being already identified in previous reports (Batz et al., 2017; Gu et al., 2013), or sharing the same seed sequence to athropod miRNAs reported in miRBase version 21. miRNAs that did not fall into any previously mentioned categories were defined as novel. By examining genomic coordinates of both lncRNAs and miRNAs, we found that 8 and 14 precursors miRNAs were fully overlapped with lncRNA loci on the same and opposite strand respectively (**Supplemental Table 3**). We classified these lncRNAs of having potential to be processed into functional mature miRNAs.

It was reported that *Ae. albopictus* was capable of producing piRNAs that were derived from protein-coding genes, suggesting that, beside TE silencing, piRNAs may possess other roles in biological pathways (Liu et al., 2016). Here, we extended the exploration of gene-derived piRNA by focusing on the piRNAs deriving from lncRNA loci. To explore this possibility, we aligned small RNA reads of 24-32 nt in size against lncRNA transcripts, and checked for the presence of typical piRNA characteristics such as 5’U bias and ping-pong signature, both of which represent hallmarks of piRNA characteristics conserved across all animal kingdom. Interestingly, we discovered that small RNA reads that aligned to lncRNAs displayed 5’U bias and ping-pong signature (**Figure 3**). The finding suggests that the biogenesis of lncRNA-derived piRNAs in *Ae. albopictus* involved both primary and secondary pathways; thereby, highlighting the possible role of piRNAs in the regulation of lncRNA expression.

**Figure 3.**
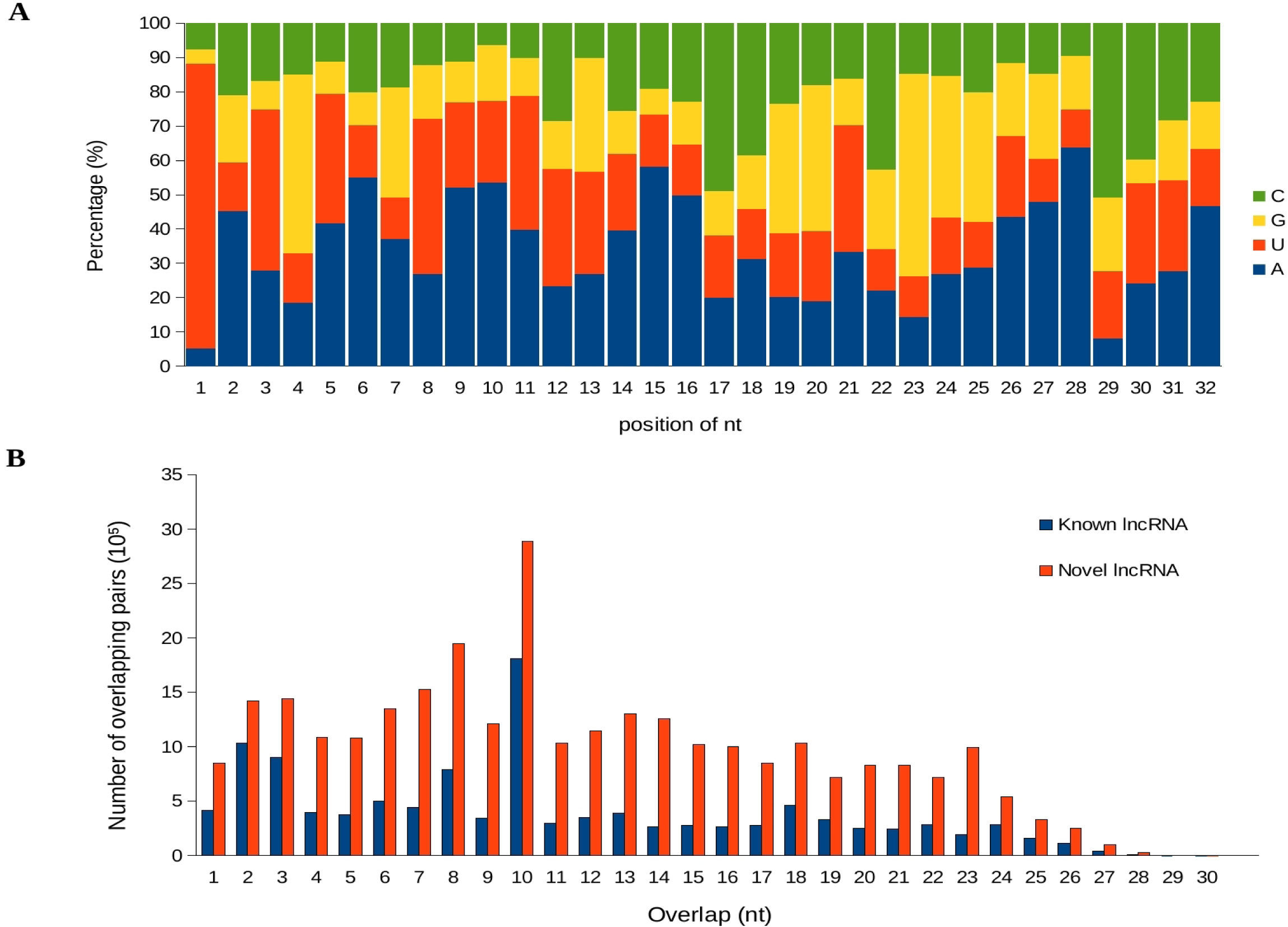
lncRNA-derived piRNA (A) Percentage of nts at each position in the reads. High percentage of U at the first position (B) Ping-pong signature of known and novel lncRNA

piRNAs were mostly transcribed from genomic loci termed piRNA clusters, which are the sources of most piRNAs (Azlan et al., 2016; Siomi et al., 2011). Since our findings pointed out the possibility of lncRNAs producing piRNAs, we then asked if lncRNA loci were largely present within piRNA clusters. To answer this question, we first identified piRNA clusters in *Ae. albopictus* genome (canu_80X_arrow2.2 assembly) using proTRAC (Rosenkranz and Zischler, 2012), and subsequently discovered that the genome harbored a total of 385 clusters (**Supplemental Table 4**). Our analysis revealed that the number of lncRNA transcripts intersecting with piRNA cluster loci was only 290, of which 160 of them were found to be fully overlapped with the clusters. The low number observed here implies that, the ability of lncRNAs serving as precusor for piRNA biogenesis is not directly due to their genomic location being present within piRNA clusters.

### Developmental expression of *Ae. albopictus* lncRNAs

In other species, lncRNAs showed high tendency to be expressed in a development-specific fashion (Cabili et al., 2011; Chen et al., 2016; Nam and Bartel, 2012; Andrea Pauli et al., 2012). To investigate if this was also true in *Ae. albopictus* lncRNAs, we analyzed public dataset (accession: SRP055126) that provided transcriptome of seven developmental stages of *Ae. albopictus* that include 0-24 and 24-48 embryonic stages, L1-L2 and L3-L4 larvae, pupae, adult males, and adult females. Consistent with findings reported in other species, across seven developmental stages, we observed that the overall expression of lncRNA was lower than protein-coding genes (**Figure 4A**).

**Figure 4.**
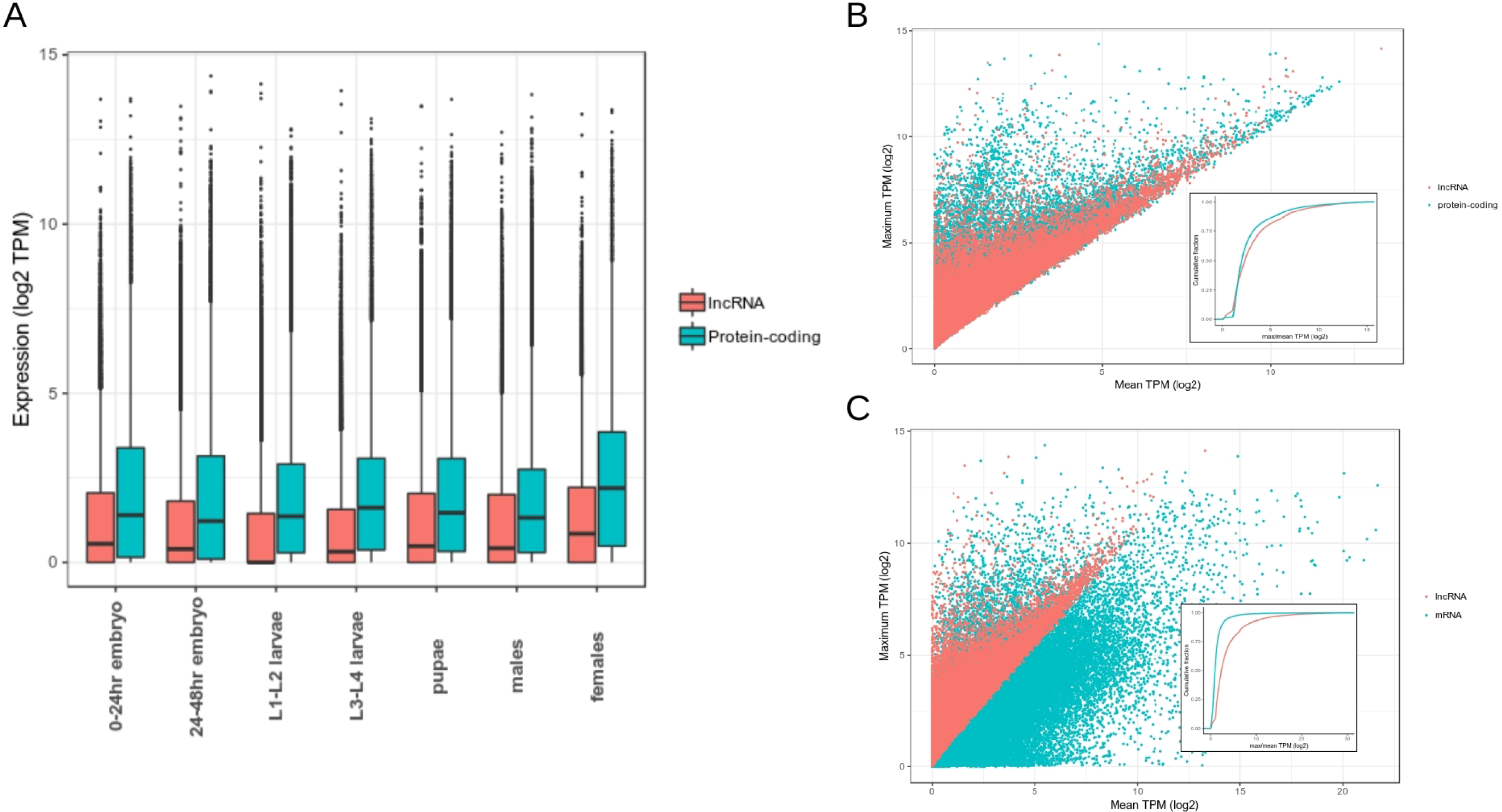
Developmental expression of *Ae. albopictus* lncRNA (A) Distribution of expression of lncRNA and protein-coding genes across seven developmental stages. (B) Differential expression at gene level of lncRNAs and protein-coding gene (C) Differential expression at transcript level of lncRNA transcripts and mRNA transcripts. In (B) and (C), for each lncRNA and protein-coding gene or mRNA, the maximum TPM value across seven developmental stages was plotted with respect to the mean of the remaining six stages. The insets within (B) and (C) display cumulative distributions of log2-scaled ratios of maximum and mean TPM of lncRNA and protein-coding gene or mRNAs.

To investigate the specificity of lncRNA expression, we compared for each gene the maximum expression among 7 developmental stages to the mean expression over the remaining 6 stages (Nam and Bartel, 2012). We repeated the same method with transcript-level expression of both lncRNA and protein-coding transcripts. By this metric, at transcript-level expression, lncRNAs were found to be more differentially expressed than mRNAs. Median fold difference between maximum and mean TPMs of lncRNAs and mRNAs were 2.4 and 0.96 respectively. On the contrary, at gene-level transcription, such difference was not observed, as median fold difference between maximum and mean TPMs of lncRNAs and coding genes were relatively similar – lncRNA was 2.4 and coding gene was 2.1.

To investigate the coexpression of lncRNAs in specific developmental stages, we conducted a hierarchical clustering analysis in Morpheus (https://software.broadinstitute.org/morpheus) based on Pearson correlation of z scores of each lncRNA (**Figure 5A**). We discovered that, compared to protein-coding genes, vast majority of lncRNAs were more tightly clustered according to specific developmental stages. We further computed specificity score of each lncRNA using an entropy-based metric of Jensen-Shannon (JS) divergence as previously described (Cabili et al., 2011). The score ranges from zero for ubiquitously expressed genes, to one for genes specifically expressed in only one tissue. Based on this measure, lncRNAs displayed more specifity as 61% of them scored 1 while the fraction of protein-coding genes showing JS score of 1 was only 28%. Therefore, the results obtained here further corroborate previous findings that claimed lncRNAs in many species displayed higher tissue or stage-specific expression compared to that of protein-coding genes (Cabili et al., 2011; A. Pauli et al., 2012; Wu et al., 2016).

**Figure 5.**
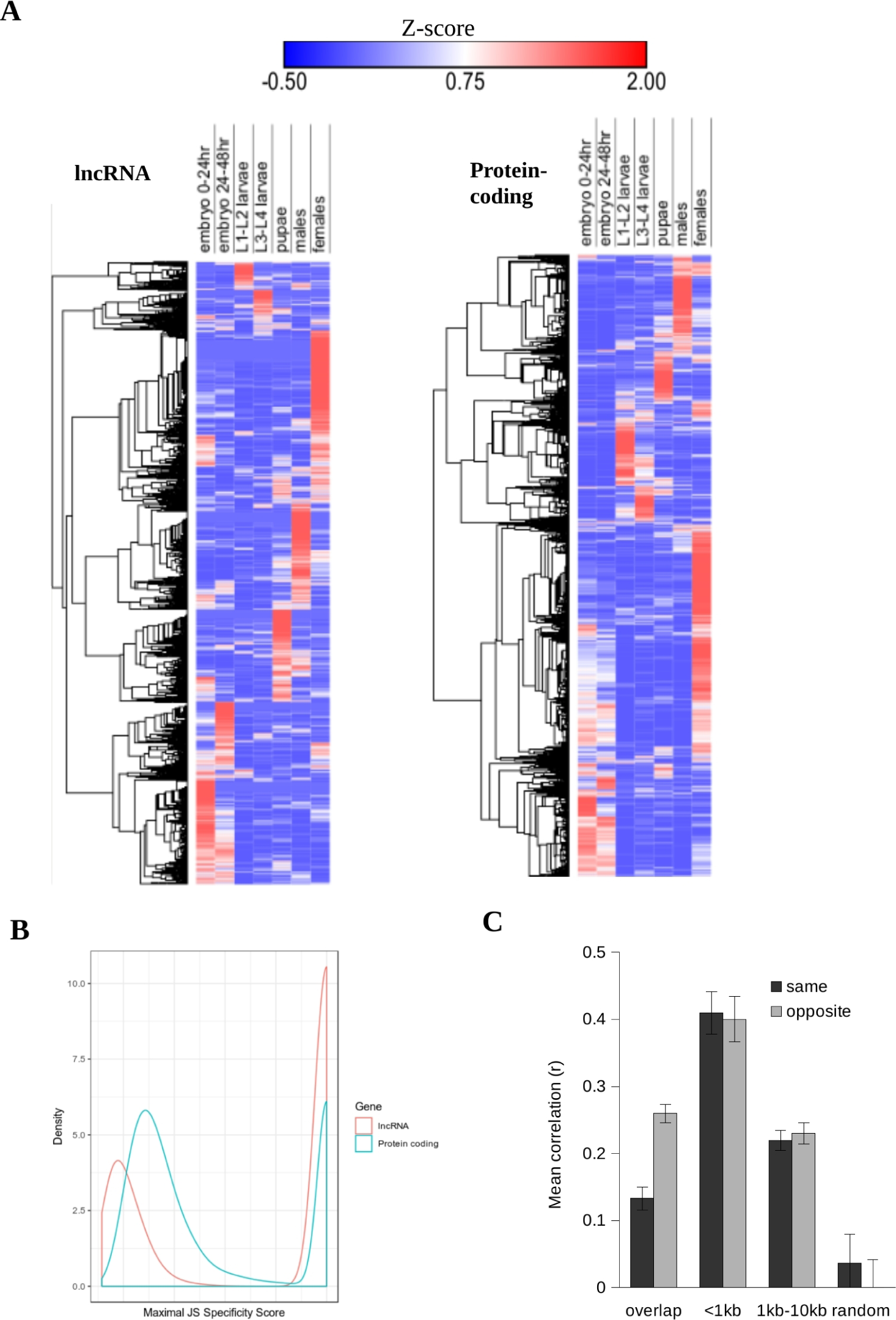
Stage-specific expression lncRNA and correlation with protein-coding gene (A) Hierarchical clustering of lncRNA and protein-coding gene across seven developmental stages. lncRNA shows tighter clustering than protein-coding gene as displayed by distinct seven nodes on the row dendogram. (B) The distribution of maximal JS specificity score across seven stages. (C) Correlation between lncRNA expression and their closest neighboring protein-coding gene. The plot shows mean Pearson correlation for lncRNA - protein-coding gene pairs in either on the same of opposite strand. Error bars show the 95% confident interval.

Previous reports showed that two neighboring genes have higher tendency to be coexpressed; thereby, showing strong correlation in expression levels (Cabili et al., 2011; Nam and Bartel, 2012; Ulitsky et al., 2011). We asked if lncRNAs in *Ae. albopictus* displayed correlation in expression between nearby or overlapping protein-coding genes that were located either on the same or opposite strand in the genome. We found that expression of the lncRNA and nearest protein-coding gene was correlated especially those within 1kb from each other, either on the same or opposite strand (Mean correlation of 0.4). As the distance increased up to 10kb, mean correlation between lncRNA and protein-coding genes lowered to around 0.2 (**Figure 5C**). Overlapping genes, on the other hand, showed less mean correlation than that of neighboring genes. Hence, this study showed that neigboring genes tended to show higher degree of expression correlation with lncRNAs than randomly assigned gene pairs

### lncRNAs were differentially expressed upon Flavivirus infection

Studies on *Ae. aegypti* transcriptomes provided the evidence that lncRNAs could be involved in DENV and ZIKV-mosquito interaction. To examine if this was also the case in *Ae. albopictus*, we generated paired-end RNA-seq libraries derived from triplicates of C6/36 cells, larval-derived *Ae. albopictus* cell line, infected with dengue virus serotype 1 (DENV1) and ZIKV. We combined both protein-coding genes and lncRNAs in our differential expression analysis using Salmon v0.9 (Patro et al., 2017) followed by edgeR (Robinson et al., 2010a). In general, we found that C6/36 cell transcriptome was highly responsive to both DENV1 and ZIKV infection. A total of 3,349 and 4,246 genes were upregulated and downregulated respectively (|log2 FC| > 1, FDR < 0.01) upon DENV1 infection (**Supplemental Table 6**). Of these genes, 1,360 and 379 of them were lncRNAs that were respectively upregulated and downregulated. Meanwhile, analysis of ZIKV-infected transcriptomes revealed a total of 3,677 upregulated genes (|log2 FC| > 1, FDR < 0.01), 1,115 of them were lncRNAs. We detected 3,979 genes (2,698 were lncRNAs) were downregulated upon ZIKV infection in C6/36 cells.

Distribution of fold change showed that upon DENV1 infection, protein-coding gene and lncRNA experienced somewhat similar level of differential expression (**Figure 6A**). Mean fold change of protein coding gene and lncRNA following DENV infection was 1.71 and 1.70 respectively. Meanwhile, in ZIKV-infected transcriptome, we discovered that mean fold change of lncRNA was higher than protein-coding gene (**Figure 6A**). For instance, lncRNA average fold change was 3.26 while protein-coding was 1.92 upon ZIKV infection. The discrepancy observed here suggest that different virus may elicit different transcriptional responses to the host cells. Besides, the observation that lncRNAs had different transcriptional response than protein-coding gene especially in ZIKV transcriptome raised the possibility that differential exression of lncRNAs was not necessarily due to the co-expression with their neighboring protein-coding genes.

**Figure 6.**
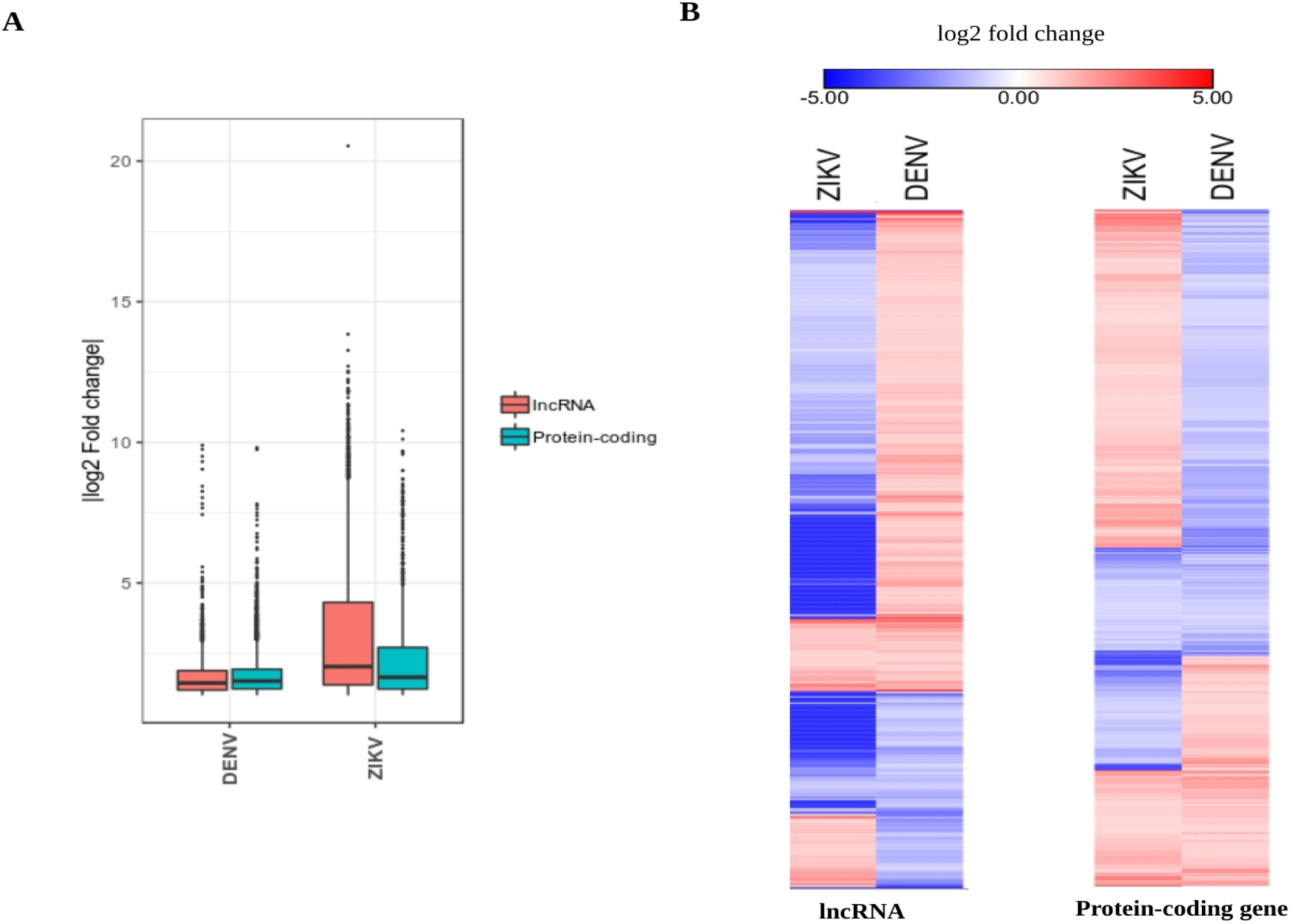
Differential expression of lncRNA upon DENV1 and ZIKV infection (A) Distribution of log2 fold change (FDR < 0.01) of lncRNA and protein-coding gene in DENV and ZIKV-infected transcriptomes. (B) Heatmaps showing the value of log2 fold change (FDR < 0.01) of lncRNA and protein-coding genes that were differentially expressed in both DENV1 and ZIKV-infected transcriptomes. Large number of genes of both lncRNA and protein-coding were found to be upregulated in one transcriptome but downregulated in another.

To investigate this possibility, we looked for protein-coding genes located closest to differentially expressed lncRNA loci, and examined how many of them having signifcant differential expression. We discovered that the number of differentially expressed protein-coding genes residing in close proximity to differentially expressed lncRNA was low. For example, in DENV transcriptome, only 225 out of 5,858 differentially expressed protein-coding genes were located closely to differentially expressed lncRNA, while in ZIKV, the number was 205 out of 3,803. Therefore, the findings further subtantiate the idea that change in lncRNA transcriptional expression landscape following virus infection was not just a byproduct of the neighboring protein-coding genes.

We detected a total of 2,071 genes that were differentially expressed in both DENV and ZIKV-infected cell transcriptomes. Of 2,071 genes, 717 of them were lncRNAs. We noticed that, compared to 2,071 genes, the number of genes that were both upregulated and downregulated in ZIKV and DENV were less than the number of genes having differential expression in opposite direction. Total number of lncRNA genes having opposite fold change in DENV and ZIKV infected transcriptomes was 506, while for protein-coding genes, the number was 905. This further support the notion that different virus evokes different transcriptional responses within the same host. Flavivirus infected transcriptome data generated in this study was limited to *Ae. albopictus* C6/36 cell line. Thus, to investigate whether the findings reported here were also true in adult *Ae. albopictus* mosquitoes, we analyzed available RNA-seq dataset on *Ae. albopictus* infected with DENV (Tsujimoto et al., 2017), and the results showed that the overall pattern of transcriptional response especially lncRNAs upon virus infection was similar regardless of tissue or cell types (**Supplemental Figure 3**).

We realized that our virus-infected transcriptomes were valuable for identifying important pathways involved in flavivirus infection. Because our main focus was to study lncRNAs, pathways and gene ontology analysis of differentially expressed protein-coding genes were not discussed here. Gene ontology analysis can be found in **Supplemental Figure 4**.

## Discussion

*Ae. albopictus,* a vector of several viruses such as DENV, ZIKV, and CHIKV, is a highly invasive species that thrives in temperate and tropical regions (Chen et al., 2015; Paupy et al., 2009). Genomic and transcriptomic investigation of *Ae. albopictus* should provide valuable genetic resources that will inform the biology of this mosquito especially on its competency as a successful vector. With this in mind, we performed global *de novo* annotation of lncRNA using RNA-seq data generated in this study and publicly available datasets. In this study, we have generated the first systematic annotation of *Ae. albopictus* transcriptome focusing primarily on lncRNAs. Huge number of sequencing reads with deep coverage enabled us to reconstruct high-confidence 23,899 lncRNA transcripts in *Ae. albopictus*. In parallel to its genome being the largest mosquito genome sequenced to date, to our knowledge, the number *Ae. albopictus* lncRNAs annotated in this study is the largest compared to other insects such as *Ae. aegypti, D. melanogaster*, and *An. gambiae* (Chen et al., 2015; Miller et al., 2018). Large genome size feature enables *Ae. albopictus* to harbor great numbers of lncRNA genes, which possibly contribute genetic materials for successful adaptation following selection in new environments (Chen et al., 2015).

Currently, three versions of *Ae. albopictus* genomes have been released – Foshan strain, AaloF1 (Chen et al., 2015), Rimini strain, AalbR1 (Dritsou et al., 2015), and C6/36, canu_80X_arrow2.2 (Miller et al., 2018). Due to small contigs and limited gene annotation of AalbR1 assembly, we only considered to use either canu_80X_arrow2.2 or AaloF1 assembly in this study. AaloF11 assembly was generated using Illumina platform while canu_80X_arrow2.2 was sequenced using PacBio technology. Due to deep coverage of long-read sequencing by PacBio technology, canu_80X_arrow2.2 assembly has larger contigs than any previously assembled mosquito genome (Miller et al., 2018); hence, offering advantage of identifying complete gene sequences. For that reason, we finally decided to use canu_80X_arrow2.2 assembly for lncRNA annotation.

Analysis of the characteristics of lncRNA in *Ae. albopictus* revealed that, despite having low level of sequence conservation among closely related insect species, lncRNAs shared strikingly similar genomic features with other species including invertebrates and vertebrates. *Ae. albopictus* lncRNAs shared many of the characteristics of their vertebrates counterparts (Hezroni et al., 2015; Andrea Pauli et al., 2012): short transcript length, relatively low expression, low level of sequence conservation, low number of isoforms, higher proportion of repeat-embedded nucleotides, low GC contents, and tend to be coexpressed with neighboring protein-coding genes. Developmental profile of transcript-level expression showed that lncRNA transcripts were more differentially expressed than mRNAs. However, this was not the case when the same analysis was performed at gene-level expression. This observation highlights the possibility that *Ae. albopictus* lncRNAs undergo active alternative splicing throughout the development, and certain isoforms are required in specific developmental stages.

Besides, lncRNAs are expressed in developmental-specific manner, and the degree of specificity is much higher than protein-coding genes. The association of specific sets of lncRNAs with well-defined developmental stage and sex, suggest that *Ae. albopictus* lncRNAs possess various roles in development. In addition, we observed that lncRNAs expressed during 0-24 hour and 24-48 hour embryo were clustered closer together, suggesting that these early embryonic lncRNAs might regulate same set of functions in embryogenesis. We also noticed that lncRNAs might have the potential to act as precursors to mature miRNAs and piRNAs, suggesting that lncRNAs are accessible to AGO/PIWI and other proteins responsible for small RNA biogenesis. Interestingly, the discovery that piRNA deriving from lncRNA transcripts was among the first to be documented. Based on the piRNA hallmarks features (5’U bias and ping-pong signature) found specifically in lncRNA-derived piRNAs, we proposed that piRNAs in *Ae. albopictus,* besides TE silencing, they might be involved in the regulation of lncRNAs.

This study also showed that genome-wide expression of lncRNAs were altered upon DENV and ZIKV infection. In line with previous studies done in mammale and *Ae. aegypti* (Etebari et al., 2017, 2016; Zhao et al., 2018), alteration of lncRNA expression landscape following flavivirus infection observed in this study implicates the involvement of *Ae. albopictus* lncRNAs in virus-host interaction. Besides, our virus-infected transcriptome analysis also revealed that, similar to other well-studied organisms (Batut and Gingeras, 2017; Engreitz et al., 2016), *Ae. albopictus* lncRNAs presumably possess their own regulatory elements that specifically govern their expression in response to viral-infection, independent of the neighboring protein-coding genes.

In summary, our study provides the first comprehensive catalog of *Ae. albopictus* lncRNA. This study also offer glimpse into lncRNA functions in numerous processes including development and virus-host response. Results generated in this study provides high-quality resources for future investigation on lncRNA functions in mosquito vectors.

## Materials and Methods

### Cell culture and virus

*Ae. albopictus* C6/36 cell (ATCC: CRL-1660) were cultured in Leibovitz’s L-15 medium (Gibco, 41300039), supplemented with 10% Fetal Bovine Serum (FBS, Gibco, 10270) and 10% Tryptose Phosphate Broth (TPB) solution (Sigma, T9157). C6/36 cells were incubated at 25°C without CO_2_. BHK-21 cells (ATCC: CCL-10) were cultured at 37°C in Dulbecco’s modified Eagles Medium (DMEM, Gibco, 11995065) supplemented with 10% FBS (Gibco, 10270), and 5% CO_2_. Dengue virus serotype 1 (Hawaiian strain) and Zika virus (Strain H/PF/2013), were propagated in C6/36 cells and titered using BHK-21 cells. Determination of DENV1 titer was done using 50% tissue culture infectious dose – cytopathic effect (TCID50-CPE) as previously described (Li et al., 2011; Atieh et al., 2016). DENV1 used in this study was a gift from Dr. David Perera, University Malaysia Sarawak. ZIKV used in this study was a gift from Dr Shee Mei Lok, Duke-NUS Medical School, Singapore.

### Virus infection, RNA extraction and sequencing

C6/36 cells were infected with DENV1 and ZIKV at multiplicity of infection (MOI) of 0.25. After 3 day post infection, RNA extraction was carried out using miRNeasy Mini Kit 50 (Qiagen, 217004) according to the manufacturer’s protocol. Total RNA was then subjected to next-generation sequencing. The RNA-sequencing libraries were prepared using standard Illumina protocols and sequenced using HiSeq2500 platform generating paired-end reads of 150 in size.

### Preparation of public datasets

Publicly available long RNA-seq datasets were downloaded from NCBI Sequence Reads Archive (SRA). List of public datasets can be found in **Supplemental Table 1**. Prior to downstream analyses, RNA-seq adapters were clipped using Trimmomatic version 0.38 (Bolger et al., 2014), and low quality reads were removed.

### Identification of lncRNA

RNA-seq libraries were mapped against *Ae. albopictus* genome (assembly: canu_80X_arrow2.2, strain: C6/36, VectorBase) using HISAT2 version 2.1.0 (Kim et al., 2015). Stringtie version 1.3.2 (Pertea et al., 2015) was used to assemble transcript, allowing the assembly of potential novel transcripts. A minimum of 200 bp size was set for transcript assembly. The resulting gtf files were merged into a using Stringtie merge, and we only retained transcripts having FPKM and RPKM of more than 0.5. Gffcompare (https://github.com/gpertea/gffcompare) was used to annotate and compare novel transcripts with the reference annotation. Transcripts with class code “i”, “u”, and “x” were retained for downstream analysis. We performed initial filtering of transcripts having coding potential using TransDecoder (Haas et al., 2013). We then evaluated further the coding potential of the remaining transcripts using CPAT (Wang et al., 2013), CPC2 (Kang et al., 2017), and CNCI (Sun et al., 2013). We applied cut-off of less than 0.3 for CPAT, and less than 0 for both CPC2, and CNCI. Only transcripts that passed the cut-off of all three softwares were retained. To exclude false positive prediction, we used BLASTX against Swissprot database, and transcripts having E-value of less than 10^-5^ were removed. We then discarded transcripts without genomic strand information.

### Differential expression and functional annotation

Salmon version Salmon v0.9 was used to quantify gene expession (Patro et al., 2017). Differential expression analysis was done using edgeR (Robinson et al., 2010b) in R/Bioconductor environment. Functional annotation was done using DAVID 6.8 (Huang et al., 2009a, 2009b) Briefly, differentially expressed transcripts were ‘BLASTX’ed against *Ae. aegypti* peptide reference (Vectorbase) with parameter of E-value < 10^-3^. The *Ae. aegypti* gene IDs were used as input in DAVID 6.8.

### miRNA identification

Analysis of miRNA discovery and expression levels were performed using the miRDeep2 v2.0.0.8 (Friedländer et al., 2012). Precursor miRNAs within the arthropod family (retrieved from miRBase version 21) were used as sequence templates of the related species (Miesen et al., 2016a). After prediction, we set two thresholds: (1) a significant Randfold p-value (p-value < 0.05) predicted Randfold v2.0.1 (Bonnet et al., 2004) for precursor miRNAs, and (2) the lowest miRDeep2 score cutoff (4.0) that had highest signal-to-noise ratio (4.5) (Ikeda et al., 2015) Predicted miRNAs that share the same homology and seed sequence with annotated miRNAs in miRBase (version 21) were categorized as homologous miRNAs, while those that do not share any sequence homology were putatively novel miRNAs.

### piRNA identification

Reads mapped to known ncRNA (snoRNAs, rRNAs, tRNAs, miscRNAs, and previously identified miRNAs) in *Ae. albopictus* were removed. Sequence of non-coding RNAs (except miRNAs) were curated from NCBI and VectorBase. The unaligned reads were considered as unannotated, and they were filtered to 24-34 nt in size. Filtered reads were then checked for the presence of 5’U bias and ping-pong signature. Ping-pong signature and 5’U bias were analyzed using perl script provided in NGS toolbox (Rosenkranz et al., 2015). Prediction of piRNA clusters was performed using proTRAC v2.4 with default settings (Rosenkranz and Zischler, 2012).

## Acknowledgements

We would like to thank all our collaborators and colleagues for the discussion and the work conducted in this lab. This study was funded by the USM Research University Grant (1001/PBIOLOGI/811320) and Sciencefund (305/PBIOLOGI/613238).

---

**Supplemental Data 1** lncRNA annotation in GTF format

**Supplemental Table 1**List of accession of public datasets used in this study

**Supplemental Table 2** Genomic coordinates of miRNA identified in this study

**Supplemental Table 3** lncRNA overlapping miRNA precurors

**Supplemental Table 4** piRNA clusters

**Supplemental Table 5**List of differentially expressed lncRNA and protein-coding genes in DENV1 and ZIKV-infected C6/36 cells

**Supplemental Table 6**List of differentially expressed lncRNA in midgut and carcass infected with dengue virus serotype 2 (DENV2) from Tsujimoto et al. 2017

**Supplemental Table 7** KEGG pathway analysis of the differentially expressed protein-coding genes in DENV1 and ZIKV-infected C6/36 cells

**Supplemental Figure 1.**
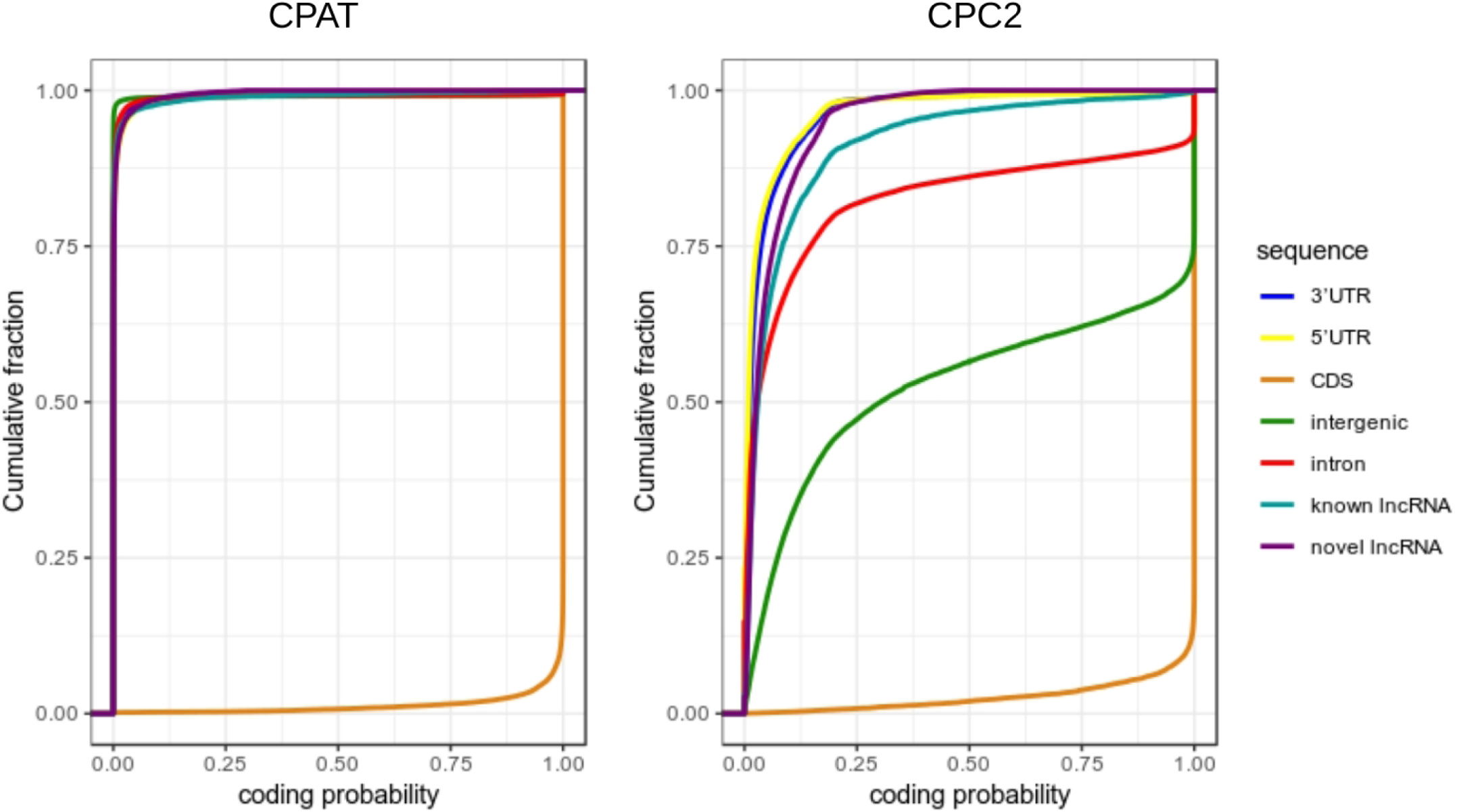
Coding probability computed by CPAT and CPC2.

**Supplemental Figure 2.**
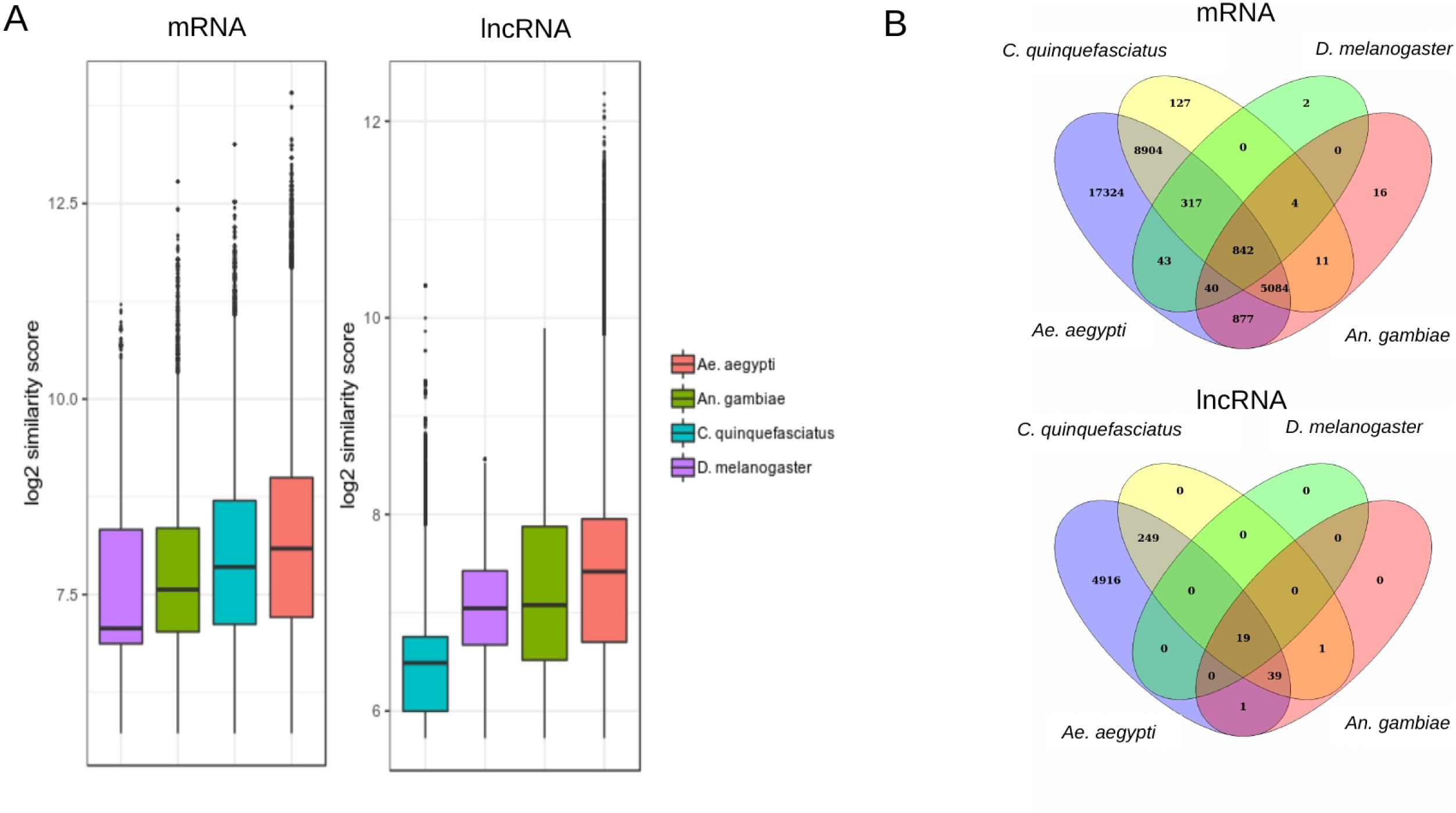
Conservation of *Ae. albopictus* lncRNA (A) Similarity bit score of *Ae. albopictus* lncRNA and mRNA transcripts with closely related insect genomes. (B) The Venn diagrams show the number of conserved *Ae. albopictus* lncRNAs and mRNAs (BLASTN e-value < 10^-50^) with closely related insect genomes.

**Supplemental Figure 3.**
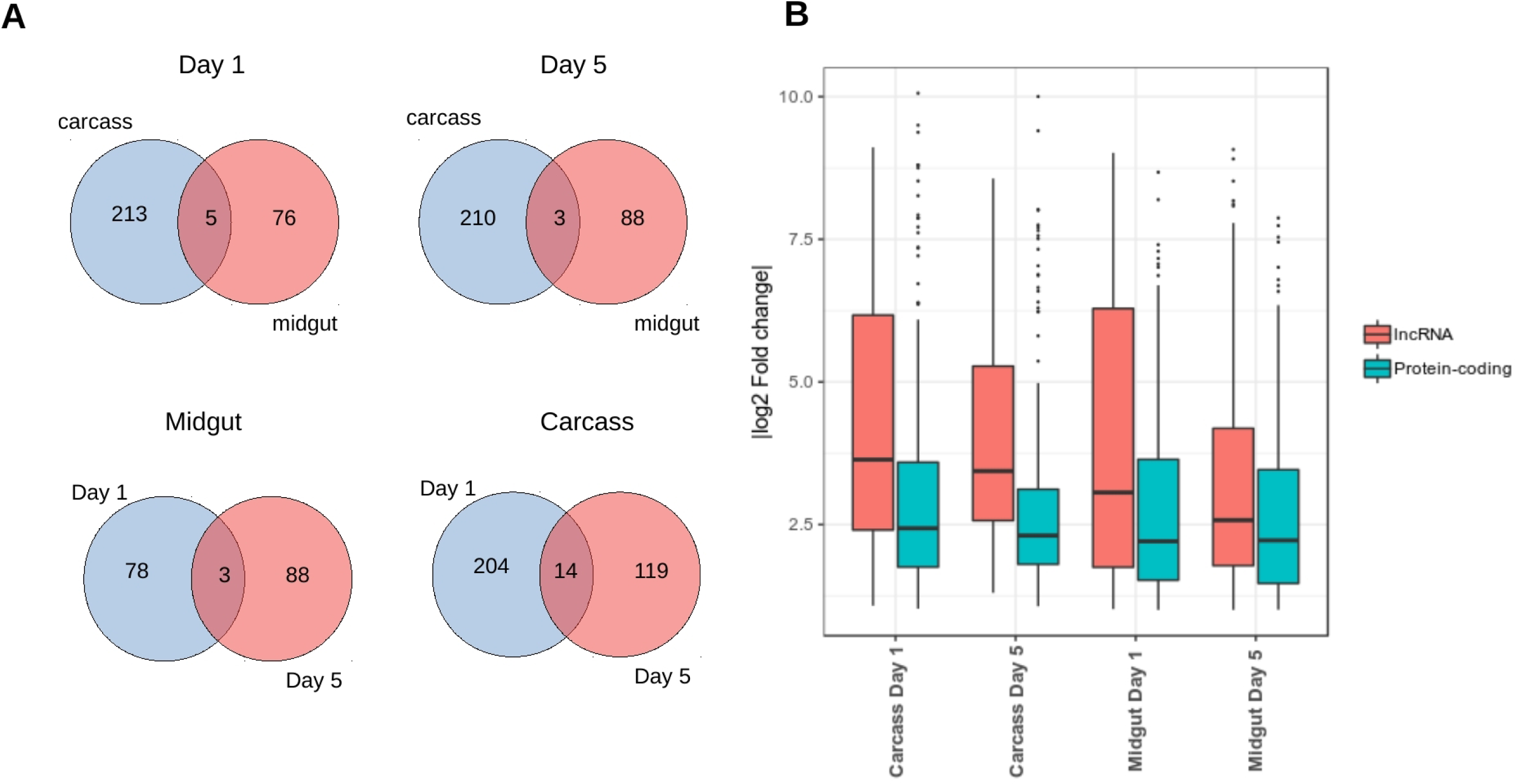
Differential expression of lncRNAs in midgut and carcass infected with dengue virus serotype 2 (DENV2) from Tsujimoto et al. 2017 (A) Number of lncRNAs that were differentially expressed (FDR < 0.01) in both carcass and midgut in day1 and day 5 post infection. (B) Distribution of log2 fold change (FDR < 0.01) of lncRNA and protein-coding gene.

**Supplemental Figure 4.**
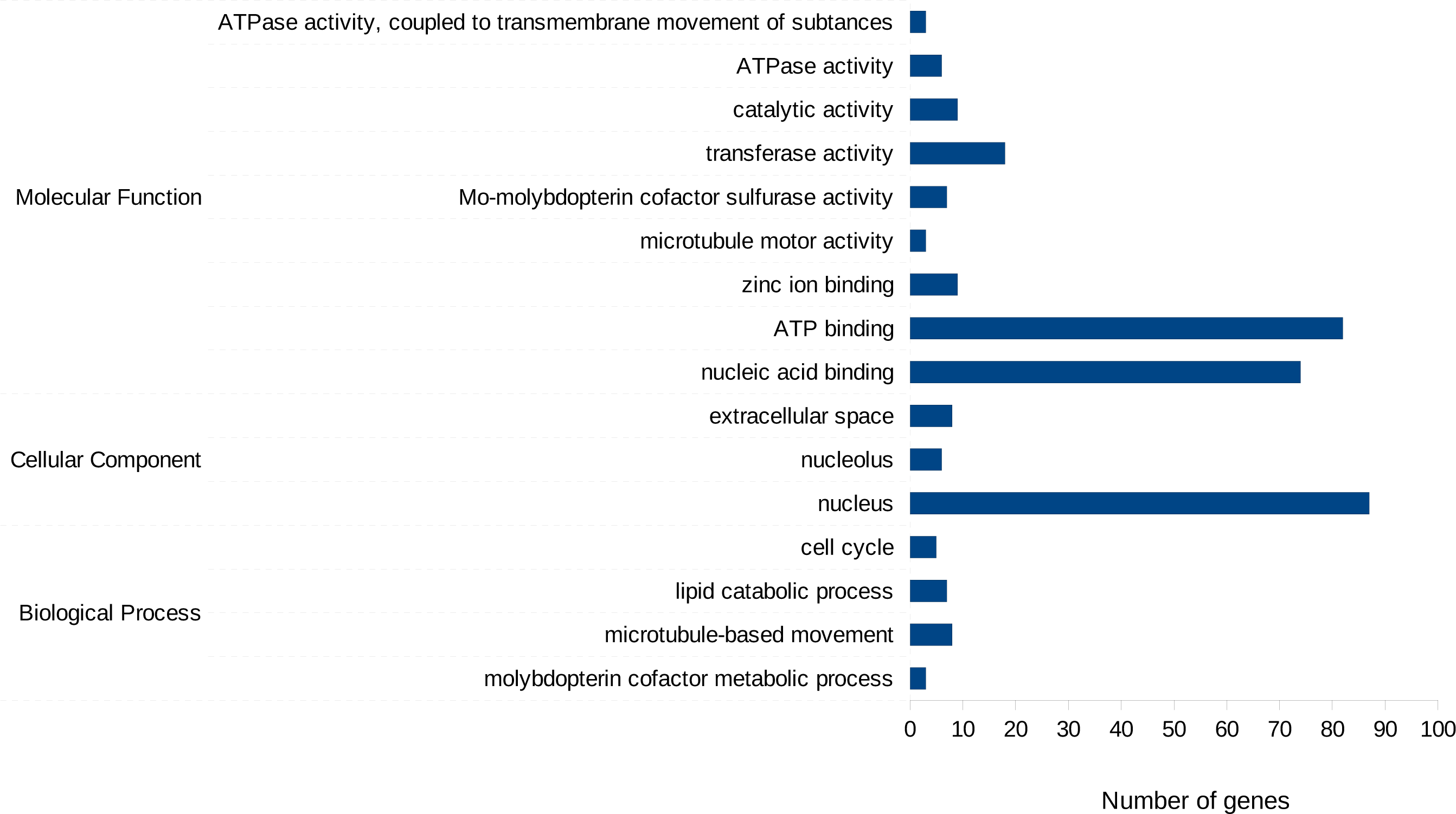
Gene ontology analysis of protein-coding genes that were differentially expressed in DENV1 and ZIKV-infected C6/36 cells.

